# *EIF4EBP1* is transcriptionally upregulated by MYCN and associates with poor prognosis in neuroblastoma

**DOI:** 10.1101/2021.12.08.471784

**Authors:** Kai Voeltzke, Katerina Scharov, Cornelius Funk, Alisa Kahler, Daniel Picard, Laura Hauffe, Martin F. Orth, Marc Remke, Irene Esposito, Thomas Kirchner, Alexander Schramm, Barak Rotblat, Thomas G. P. Grünewald, Guido Reifenberger, Gabriel Leprivier

## Abstract

**Background:** Neuroblastoma (NB) accounts for 15% of cancer-related deaths in childhood despite considerable therapeutic improvements. While several risk factors, including *MYCN* amplification and alterations in RAS and p53 pathway genes, have been defined in NB, the clinical outcome is very variable and difficult to predict. Since genes of the mTOR pathway are up-regulated in *MYCN*-amplified NB, we aimed to define the predictive value of the mTOR substrate-encoding gene eukaryotic translation initiation factor 4E-binding protein 1 (*EIF4EBP1*) expression in NB patients.

**Methods:** Several independent NB patient cohorts with corresponding mRNA expression data were analyzed for *EIF4EBP1* expression. An institutional NB cohort consisting of 69 prospectively collected tumors was employed to immunohistochemically analyze expression of *EIF4EBP1*-encoded protein (4EBP1). In addition, we performed an *in vitro* luciferase reporter gene assay with an episomal *EIF4EBP1* promoter and genetically modulated *MYCN* expression in NB cells.

**Findings:** *EIF4EBP1* mRNA expression was positively correlated with *MYCN* expression and elevated in stage 4 and high-risk NB patients. High *EIF4EBP1* mRNA expression was associated with reduced overall and event-free survival in the entire group of NB patients in three cohorts, as well as in stage 4 and high-risk patients. High levels of 4EBP1 were significantly associated with prognostically unfavorable NB histology. Functional analyses *in vitro* revealed that *EIF4EBP1* expression is transcriptionally controlled by MYCN binding to the *EIF4EBP1* promoter.

**Interpretation:** High *EIF4EBP1* expression is associated with poor prognosis in NB patients and may serve to stratify patients with high-risk NB.

**Funding:** G.L. was supported by funding from the Elterninitiative Düsseldorf e.V., the Research Commission of the Medical Faculty of Heinrich Heine University, the Deutsche Forschungsgemeinschaft (Grant LE 3751/2-1), and the German Cancer Aid (Grant 70112624). The laboratory of T.G.P.G. is supported by the Barbara und Wilfried Mohr Foundation. BR is supported by the Israel Science Foundation (grant No. 1436/19).

**RESEARCH IN CONTEXT:** *Evidence before this study:* NB represents a particularly heterogeneous cancer entity, with 5-year event-free survival rate ranging from 50% to 98% depending on the patient’s risk group. While genes of the nutrient-sensing mTOR pathway were found to be up-regulated in *MYCN*-amplified NB tumors, their clinical relevance and prognostic value in NB patients remain unclear. In particular, the mTOR substrate-encoding gene *EIF4EBP1* was studied in NB by three different groups and high *EIF4EBP1* mRNA expression was observed in *MYCN*-amplified or contradictorily in more favorable stages 1 and 2 patients. Also, *EIF4EBP1* was included in a prognostic gene signature for poor overall survival in NB. However, the prognostic value of *EIF4EBP1* alone was not determined in NB and the expression of *EIF4EBP1* encoded protein, 4EBP1, was not analyzed in NB tumor tissues and not correlated with clinicopathological features such as histological subtypes. Additionally, the transcriptional regulation of the *EIF4EBP1* promoter by MYCN was not characterized.

*Added value of this study:* This study uncovers the prognostic potential of *EIF4EBP1* at the mRNA and protein levels in NB patients. We report that high *EIF4EBP1* expression is correlated with poor survival in three independent cohorts and that high 4EBP1 levels is associated with a prognostically unfavorable histological subtype. High *EIF4EBP1* expression is also a factor of poor prognosis in stage 4 and high-risk patient groups. Finally, we found that MYCN activates the human *EIF4EBP1* promoter through binding at three binding motifs.

*Implications of all the available evidence:* *EIF4EBP1* mRNA and 4EBP1 protein expression have prognostic value in NB, especially to stratify patients with advanced and more aggressive NB, such as patients with stage 4 disease and high-risk patients including those with unfavorable histological subtype NB. Enhanced *EIF4EBP1* mRNA and 4EBP1 protein expression in NB are driven by direct transcriptional activation of *EIF4EBP1* by MYCN.

## INTRODUCTION

Neuroblastoma (NB) is a pediatric malignant tumor that develops from progenitor cells of the sympathetic nervous system and the adrenal glands [1,2]. NB is the most commonly occurring extracranial solid tumor in childhood and the major cause of cancer-related mortality in infants [2]. NB tumors are classified into five stages (1, 2, 3, 4 and 4S) according to tumor size, the presence of metastasis and the outcome of surgical resection [1]. Noteworthy, stage 4S represents a special form of NB in infants that is associated with a high chance of spontaneous regression despite metastatic spread [1]. Apart from surgical resection, treatment options may include response-adjusted chemotherapy for low to intermediate risk groups or a mix of surgery, high-dose chemotherapy, immunotherapy, and radiation for patients belonging to the high-risk group. The risk level is determined based on the tumor stage combined with age at diagnosis, tumor ploidy, genetic alterations and tumor histology [1,3]. However, NB represents a particularly heterogeneous type of cancer, posing challenges to precisely predict therapeutic response and clinical outcome in the individual patient [4,5]. While some NB tumors may spontaneously regress, high-risk patients have an increased likelihood of relapse and available treatment options for relapsed patients are rarely successful. Indeed, the 5-year event-free survival rate for high-risk patients is less than 50%, in contrast to 90-98% for low-risk patients [6]. In addition, success rates of second line treatment in relapsed patients remain poor [5,7]. Therefore, it is critical to define novel stratification factors for NB patients to better predict individual risk and to facilitate administration of the most appropriate therapeutic option.

NB is rarely familial (1-2%) and only few predisposition genes, such as *PHOX2B* and *ALK*, have been reported [4,8–10]. Genetically, several acquired alterations have been detected in NB and linked to patient outcome. These include gain-of-function mutations in *ALK*, gain of chromosome arm 17q, loss of chromosome arm 11q, amplification of *MYCN* [4], and, more recently reported, alterations in genes related to the RAS and p53 pathways [11]. *MYCN* amplification is found in about 20% of NB and is associated with aggressive tumors, therapy resistance and poor survival [6]. *MYCN* is a member of the *MYC* oncogene family and encodes a transcription factor that recognizes a specific DNA element referred to as E-box [12,13]. This allows MYCN to regulate the transcription of genes involved in cell cycle progression, proliferation, differentiation and survival [6]. MYCN is a strong driver of NB tumorigenesis, as tissue-specific overexpression of MYCN is sufficient to induce NB tumor development in mouse models [14]. Mechanistically, MYCN is proposed to rewire metabolism to enable NB tumor cells to proliferate, in turn preserving the intracellular redox balance while producing enough energy by inducing a glycolytic switch [15–17]. In particular, MYCN actively augments the transcription of multiple genes whose products are involved in the protein synthesis machinery [16]. Even though MYCN represents a highly attractive therapeutic target in NB, as a transcription factor that lacks hydrophobic pockets which can be targeted by drug-like small molecules, it is still considered as being “undruggable” [18,19]. Thus, identification of downstream effectors involved in MYCN-driven NB progression is a promising approach to uncover novel targets for molecularly guided therapeutic approaches. To better delineate the molecular basis of *MYCN*-amplified NB aggressiveness, several approaches have been undertaken. In particular, RNA-sequencing (RNA-seq) has been used to uncover the set of genes induced in *MYCN*-amplified compared to *MYCN*-non-amplified NB [20]. Strikingly, this analysis identified regulators of protein synthesis which are components of the mechanistic target of rapamycin (mTOR) pathway, including the mTOR target eukaryotic initiation factor 4E binding protein 1 (*EIF4EBP1*). The corresponding protein, 4EBP1, is inhibited through mTOR-mediated phosphorylation when nutrients are available, leading to active mRNA translation initiation [21]. Under nutrient-deprived conditions, when mTOR is inhibited, 4EBP1 gets activated and thus binds to the translation initiation factor eIF4E, in turn blocking cap-dependent mRNA translation initiation [21]. At the cellular level, 4EBP1 is negatively regulating proliferation and mitochondrial activity [22,23]. The exact role of 4EBP1 in cancer is still debated. 4EBP1 was found to exert a tumor suppressive function *in vivo*, as 4EBP1 knock-out leads to enhanced tumor formation in mouse models of head and neck squamous cell carcinoma [24], and prostate cancer [25]. In contrast, 4EBP1 was shown to mediate angiogenesis and facilitate tumor growth in a breast cancer model *in vivo*, highlighting a cancer type-specific function of 4EBP1 [26]. In keeping with that, the clinical relevance of *EIF4EBP1* expression depends on the tumor type.

*EIF4EBP1* was reported to be overexpressed in a number of tumor entities in adults [27], including breast cancer [28], in which *EIF4EBP1* is amplified as part of the 8p11-12 amplicon, as well as in ovarian and prostate cancer [29,30]. In breast and liver cancer, high *EIF4EBP1* expression has been associated with poor survival [28,31]. In contrast, *EIF4EBP1* expression was found to be reduced in head and neck cancer, in which low expression is correlated with poor prognosis [24]. In NB, the expression of *EIF4EBP1* is deregulated, even though contradictory findings have been reported. While *EIF4EBP1* was characterized as a gene upregulated in *MYCN*-amplified versus *MYCN*-non-amplified NB tissues and cells [20], another study reported that *EIF4EBP1* levels were higher in favorable stages of NB as compared to advanced stage 4 tumors [32]. In addition, Meng *et al.* showed that *EIF4EBP1* is part of a gene signature that predicts poor overall survival [33]. However, it was not investigated whether *EIF4EBP1* expression alone can predict NB patient prognosis. Thus, the clinical relevance of *EIF4EBP1* expression in NB needs further evaluation.

Overexpression of *EIF4EBP1* in cancer is mediated by certain transcription factors, such as MYC [34], androgen receptor [35], and the stress regulators ATF4 [36] and HIF-1α [37], which all bind to and thereby modulate the activity of the *EIF4EBP1* promoter. More specifically, ChIP-sequencing (ChIP-seq) revealed binding of MYCN to the *EIF4EBP1* promoter in NB cells, and MYCN was reported to impact *EIF4EBP1* transcription, pointing to *EIF4EBP1* as a potential MYCN target gene [38,39]. However, how MYCN exactly controls the *EIF4EBP1* promoter is still poorly understood.

In this study, we analyzed publicly available NB patient data sets and revealed that *EIF4EBP1* is overexpressed in NB compared to normal tissues, is significantly co-expressed with *MYCN*, and is elevated in high-risk relatively to low-risk tumor groups. High *EIF4EBP1* levels were found to be significantly linked to poor overall survival in all NB patients, as well as in the more aggressive stage 4 and high-risk groups. In addition, immunohistochemistry staining of NB tissues confirmed the mRNA-based associations and showed that high 4EBP1 protein expression associates with unfavorable histology in NB. Finally, by applying gene reporter assays and by modulating MYCN expression *in vitro*, we found that MYCN upregulates the *EIF4EBP1* promoter activity by binding to three distinct E-boxes.

## MATERIALS AND METHODS

### Databases

The RNA-seq, microarray and ChIP-seq data were retrieved from ‘R2: Genomics Analysis and Visualization Platform’ (http://r2.amc.nl). Data were visualized with IGV or Affinity Designer. For the MYCN occupancy profile in BE(2)-C cells, the ChIP-seq data by Durbin *et al.* (GSE94824) were accessed using the human genome GRCh 38/hg 38. For the initial across dataset analysis, four publicly available and independent cohorts, namely the Versteeg *et al.* (GSE16476), Lastowska *et al.* (GSE13136), Hiyama *et al.* (GSE16237), and Delattre *et al.* (GSE14880) datasets were used. The remaining expression, amplification and survival data consisted of the independent SEQC/ MAQC-III Consortium GSE49710), Kocak *et al.* study (GSE45547) and Neuroblastoma Research Consortium [NRC] (GSE49710), Kocak (GSE45547) and NRC (GSE85047) cohorts. For the expression analysis of TH-MYCN transgenic NB model, the dataset from Balamuth *et al.* (GSE17740) was used.

### Immunohistochemistry

For immunohistochemistry, deparaffinated tissue sections were pretreated with citrate buffer at 98°C for 20 min, cooled down to room temperature, and blocked with 2% horse serum, avidin blocking solution and biotin blocking solution (Avidin/Biotin Blocking Kit, SP-2001, Vector Laboratories, Burlingame, CA, USA) for 10 min each. Staining for 4EBP1 was carried out with monoclonal anti-4EBP1 raised in rabbit (1:200; ab32024, Abcam, Cambridge, UK) for 2 h at 37°C. Detection was carried out using the Dako REAL detection system, alkaline phosphatase/RED, rabbit/mouse following manufacturer’s instructions (Detection Kit #K5005, Agilent Technologies, Santa Clara, CA, USA). Immunostained tissue sections were counterstained with hematoxylin solution according to Mayer (T865.1, Roth, Karlsruhe, Germany).

Evaluation of immunoreactivity of 4EBP1 was carried out in analogy to scoring of hormone receptor Immune Reactive Score (IRS) ranging from 0–12. The percentage of cells with expression of the given antigen was scored and classified in five grades (grade 0 = 0–19%, grade 1 = 20–39%, grade 2 = 40–59%, grade 3 = 60–79% and grade 4 = 80−100%). In addition, the intensity of marker immunoreactivity was determined (grade 0 = none, grade 1 = low, grade 2 = moderate and grade 3 = strong). The product of these two grades defined the final IRS. IRS 0-6 was considered as “low” staining level while IRS 7-12 was categorized as “high” staining level.

Tissue microarrays (TMAs) were constructed by taking three representative cores (each 1 mm in diameter) from respective blocks exhibiting at least 80% viable tumor tissue. Tumor blocks were retrieved from the archives of the Institutes of Pathology of the LMU Munich or the University Hospital Düsseldorf with IRB approval (study numbers 550-16 UE for LMU Munich and 2018-174 for the University Hospital Düsseldorf).

### Statistics

All experiments were, if not otherwise stated, independently carried out at least three times. Statistical significance was calculated using Student’s t-test or Mann-Whitney U-test in GraphPad Prism 8. For survival analysis, the cohorts were stratified based on relative expression of *EIF4EBP1*. The median was chosen as expression cutoff to determine high and low *EIF4EBP1* level. Statistical significance was determined by the logrank test. Multivariate analysis was performed using the Cox Regression method in SPSS v21 (IBM). To calculate significance of the scoring of immunohistochemistry staining, the Chi-square test was used. The data are represented as means +/− standard deviation. A p-value of less than 0.05 was considered significant.

### Cell culture

Cells were maintained using standard tissue culture procedures in a humidified incubator at 37°C with 5% CO_2_ and atmospheric oxygen. NB cell lines IMR-32 and Kelly, and HEK-293-T cells were obtained from American Type Culture Collections (ATCC, Manassas, VA, USA). SHEP-TR-MYCN engineered NB cell lines have been previously described [17]. NB cell lines were cultured in Roswell Park Memorial Institute (RPMI)-1640 medium (Thermo Fisher Scientific, Waltham, MA, USA), while HEK-293-T cells were maintained in Dulbecco’s modified Eagle medium (DMEM) (Thermo Fisher Scientific). All cell culture media were supplemented with 10% (volume/volume) fetal bovine serum (FBS) (Sigma-Aldrich, St. Louis, MI, USA) and 1% penicillin/streptomycin (Thermo Fisher Scientific). Cells were treated with 3 μg/ml plasmocin (Invivogen, San Diego, CA, USA) to prevent mycoplasma contamination. To induce MYCN expression, SHEP-TR-MYCN cells were treated with 1 μg/ml doxycycline. All cell lines were routinely confirmed to be mycoplasma-free using Venor^®^GeM Classic kit (Minerva Biolabs, Berlin, Germany). Cell lines were authenticated by STR-profiling (Genomics and Transcriptomics Laboratory, Heinrich Heine University, Germany).

### RNA extraction, cDNA synthesis and quantitative real time PCR

Total RNA was purified from cells using the RNeasy plus mini kit (QIAgen, Hilden, Germany) according to the manufacturer’s handbook. RNA concentration and purity were assessed by spectrophotometry using the NanoDrop2000 (Thermo Fisher Scientific). Subsequently, each sample was diluted to a concentration of 100 ng/μl in nuclease-free water. For cDNA synthesis, 1 μg RNA was processed in a total reaction volume of 20 μl using the High-Capacity cDNA Reverse Transcription kit (Applied Biosystems, Waltham, MA, USA), following the manufacturer’s protocol. Quantitative real time reverse transcription PCR was performed using SYBR green PCR master mix (Applied Biosystems) and the CFX384 Touch Real-Time PCR Detection System (Bio-Rad Laboratories, Hercules, CA, USA). Relative expression levels of *MYCN* and *EIF4EBP1* were normalized to internal housekeeping genes *GUSB* and *PPIA*. The primer list can be found in supplementary table 1.

### Immunoblot analysis of protein expression

Cells were washed with phosphate buffered saline (PBS) and lysed in radioimmunoprecipitation assay (RIPA) buffer (150 mM NaCl, 50 mM Tris-HCl, pH 8, 1% Triton X-100, 0.5% sodium deoxycholate, and 0.1% SDS) supplemented with protease inhibitors (Sigma-Aldrich) and phosphatase inhibitors mix (PhosphoSTOP, Roche, Penzberg, Germany). Cell lysates were centrifuged at 21,000 rpm for 15 min at 4°C to separate cell debris and DNA from protein lysates. Protein concentration was measured with the BCA protein assay kit (Thermo Fisher Scientific), according to manufacturer’s protocol. Protein lysates were separated by SDS-PAGE and transferred onto a nylon membrane. The membrane was incubated for 1 h in Tris-buffered saline Tween (TBST) (50 mM Tris-Cl, 150 mM NaCl, pH 7.5, 0.1% Tween-20) containing 5% bovine serum albumin (BSA), to prevent non-specific antibody binding, followed by an overnight incubation at 4°C with the following primary antibodies: 4EBP1 (1:1,000, Cell Signaling Technology, Cambridge, UK #9644), MYCN (1:1,000, Cell Signaling #9405), GAPDH (1:1,000, Cell Signaling #2118), and β-Actin (1:5,000, Sigma-Aldrich #A2228). The secondary antibodies IRDye 800CW Goat anti-Rabbit (1:10,000, LI-COR Biosciences, Bad Homburg, Germany #926-32211) or IRDye 800CW Goat anti-Mouse (1:10,000, LI-COR Biosciences #926-32210) were incubated at room temperature for 1 h, followed by detection of the fluorescent signal with the Odyssey CLx imager (LI-COR Biosciences).

### Plasmid construction

The promoter region of the human *EIF4EBP1* gene, spanning from −192 to +1372, was inserted into the SacI and BglII restriction sites of the Firefly Luciferase expressing pGL4.22 plasmid (Promega, Madison, WI, USA). Each of the three identified MYCN binding site was subsequently mutated alone or in a combination of two sites. Each of the E-box sequence has been mutated to CAAGGC. All cloning was performed by GENEWIZ Germany GmbH (Leipzig, Germany).

### Luciferase Reporter Assay

For the promoter reporter assay, HEK-293-T cells were seeded into 12-well plates and co-transfected the following day with 500 ng of the *EIF4EBP1* WT or mutant promoter pGL4.22 plasmids, 50 ng of the MYCN overexpressing pcDNA3.1 plasmid or empty pcDNA3.1 plasmid, and 3 ng of the *Renilla* Luciferase expressing pRL-SV40 plasmid (Promega) for normalization. For transfection, plasmids were incubated with 3 μl CalFectin (SignaGen laboratories, Rockville, MD, USA) in Opti-MEM (Thermo Fisher Scientific) for 20 min before adding the mix dropwise onto the cells. 48 h post-transfection, cells were passively lysed and processed according to the protocol of the Dual-Luciferase^®^ Reporter Assay System (Promega), besides using only half the recommended volume of detection buffers. Firefly and *Renilla* luciferase activities were sequentially measured using a Tecan Spark plate reader and the ratio of firefly luciferase to *Renilla* luciferase luminescence was calculated. The experiments were repeated independently for three times.

## RESULTS

### *EIF4EBP1* expression is increased in NB and correlates with *MYCN* expression

To assess the clinical significance of *EIF4EBP1* expression, we first examined *EIF4EBP1* mRNA levels in NB tumor tissue samples and normal tissues. We pooled microarray data of four different NB cohorts and retrieved expression data from adrenal tissue used as the corresponding normal tissue (Fig. 1a). This indicated that *EIF4EBP1* expression is significantly elevated in NB compared to adrenal gland (p<0.0001, Fig. 1a). We then determined whether *EIF4EBP1* expression is related to the *MYCN* amplification status. By comparing the level of *EIF4EBP1* in *MYCN*-amplified versus *MYCN*-non-amplified NB samples, we found that *EIF4EBP1* is expressed at higher levels in *MYCN*-amplified compared to *MYCN*-non-amplified NB in the SEQC and Kocak cohorts [40,41] (p<0.0001, Fig. 1b; p<0.0001, Fig. 1c). This further supports and extends previous observations made in a limited number of NB samples (n=20) showing *EIF4EBP1* overexpression in *MYCN*-amplified versus *MYCN*-non-amplified NB tumors [20]. Since *MYCN* amplification may result in different levels of *MYCN*, we next investigated whether expression levels of *MYCN* and *EIF4EBP1* in NB correlate with each other. Our analyses highlight a significant coexpression between *MYCN* and *EIF4EBP1* in the SEQC (correlation coefficient [r]=0.564, p<0.0001, Fig. 1d) and Kocak ([r]=0.532, p<0.0001, Fig. 1e) cohorts. These findings are in line with the reports that *EIF4EBP1* is a potential MYCN target gene in NB [38,39]. We also assessed whether the expression of *EIF4EBP1* is determined by NB stages or risk groups, and found that *EIF4EBP1* levels are increased according to NB tumor aggressiveness in two cohorts (Fig. 1f&g). In particular, *EIF4EBP1* is expressed at higher levels in stage 4 NB tumors as compared to stage 1 and stage 2 tumors (stage 4 versus stage 1, p<0.0001, Fig. 1f; p<0.0001 Fig. 1g). Interestingly, samples from stage 4S NB showed significantly lower *EIF4EBP1* levels compared to stage 4 tumors (stage 4S versus stage 4, p<0.01, Fig. 1f; p<0.001, Fig. 1g). In support of this finding, we observed that in the SEQC cohort *EIF4EBP1* expression is higher in high-risk compared to low-risk NB, as based on the Children’s Oncology Group (COG) classification (p<0.0001, Fig. 1h). Such clinical information was not available in any other publicly available cohorts with mRNA expression data. Taken together, we present evidence that *EIF4EBP1* is commonly overexpressed in NB tumors and that *EIF4EBP1* level is increased in *MYCN*-amplified NB and advanced NB stages.

**Figure 1:**
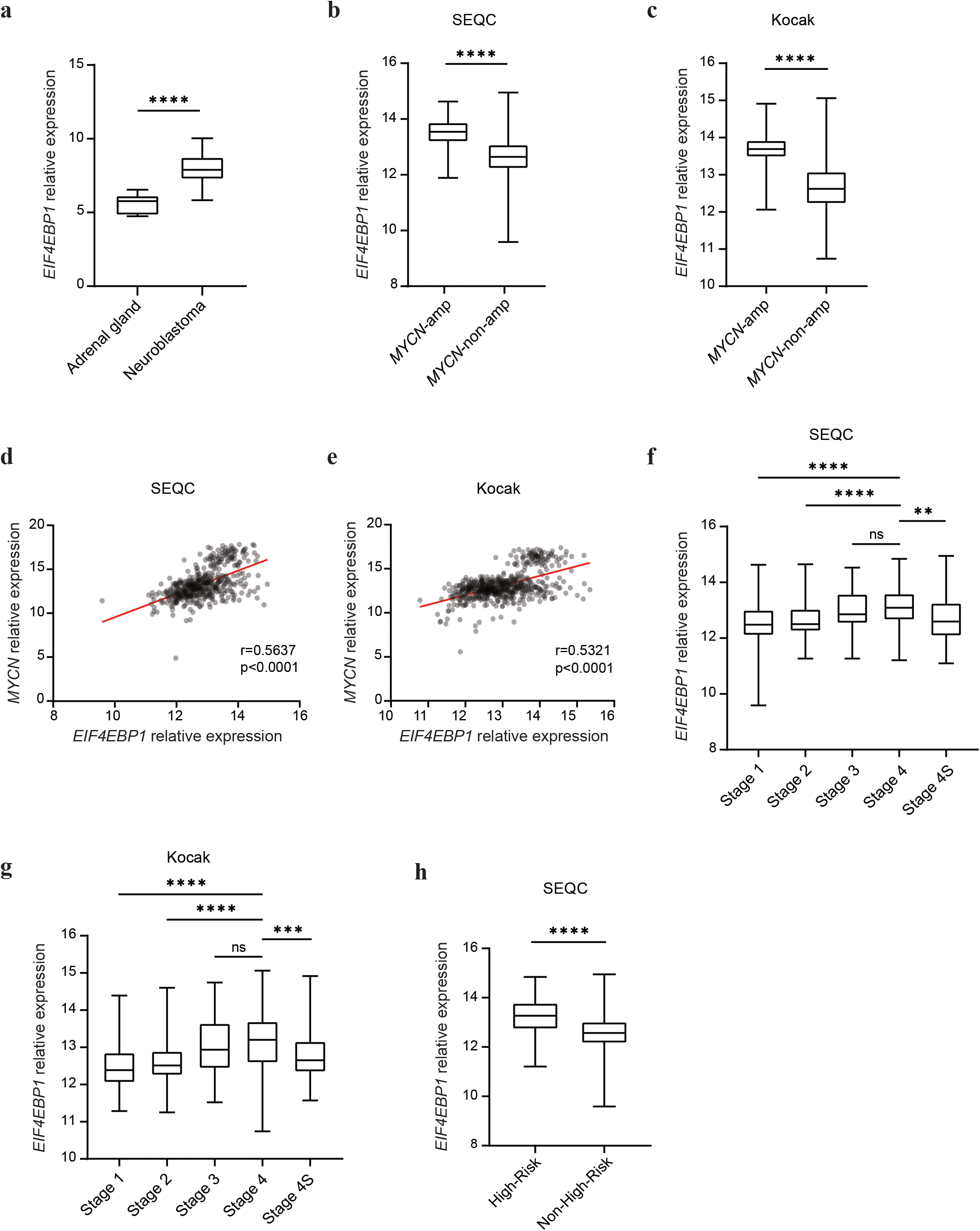
*EIF4EBP1* mRNA expression is associated with *MYCN* mRNA expression and is increased in more advanced and aggressive NB subsets. (a) Expression levels of *EIF4EBP1* mRNA in a pool of four different NB cohorts (total n=203), compared to healthy control tissues (adrenal gland, n=13). (b, c) Expression levels of *EIF4EBP1* mRNA n *MYCN*-amplified (n=92, SEQC [b] and n=93, Kocak [c]) compared to *MYCN*-non-amplified (n=401, SEQC [b] and n=550 Kocak [c]) NB patients of the SEQC (b) and Kocak (c) cohorts. (d, e) Expression levels of *EIF4EBP1* mRNA plotted against expression levels of *MYCN* mRNA in SEQC (r=0.5637, d) and Kocak (r=0.5321, e) cohorts. (f, g) Expression levels of *EIF4EBP1* mRNA per NB stage in SEQC (f) and Kocak (g) cohorts. (h) Expression levels of *EIF4EBP1* mRNA in high-risk (n=176) compared to non-high-risk (n=322) NB in the SEQC cohort. Data were retrieved from the R2: Genomics Analysis and Visualization Platform. Statistics were determined using Mann-Whitney U-test. Exact p-values are presented. \**P*<0.05, \*\**P*<0.01, \*\*\**P*<0.001, \*\*\*\**P*<0.0001.

### *EIF4EBP1* expression is a factor of poor prognosis in NB

Since we found *EIF4EBP1* mRNA levels to be elevated in aggressive NB subsets, we examined whether *EIF4EBP1* expression is linked to prognosis in NB patients. Kaplan-Meier estimates univocally showed that high *EIF4EBP1* levels (using median expression level as cut off) were significantly associated with reduced overall and event-free survival in three independent cohorts, namely SEQC, Kocak and NRC cohorts [42] (p=3.1e-08, Fig. 2a; p=4.2e-11, Fig. 2b; p=1.7e-06, Fig. 2c, and supplementary Fig. 1a, b&c). To test dependence of *EIF4EBP1* expression as prognostic factor on established factors of poor prognosis, we performed multivariate analysis to determine the statistical interaction between high *EIF4EBP1* expression and *MYCN* amplification status, tumor stage or age at diagnosis. This indicated that *MYCN* amplification status, tumor stage and age at diagnosis each influenced the prognostic value of high *EIF4EBP1* expression in the SEQC and NRC cohorts (Tables 1&2). Therefore, high *EIF4EBP1* expression is not an independent factor of poor prognosis in NB. However, we uncovered that *EIF4EBP1* expression can predict overall survival in clinically relevant NB subsets, including more advanced and aggressive NB subgroups. Indeed, our analyses highlighted that high *EIF4EBP1* expression significantly predicted reduced overall survival in *MYCN*-non-amplified patients of the SEQC and NRC cohorts (p=3.8e-03, Fig. 2d; p=0.04, Fig. 2e), while it was significant for event-free survival only in the SEQC cohort (supplementary Fig. 1d&e). On the other hand, Kaplan-Meier survival estimates in high-risk NB patients (SEQC cohort) revealed that high *EIF4EBP1* levels were correlated with poor overall survival (p=7.4e-03, Fig. 2f), as well as with reduced event-free survival (supplementary Fig. 1f), suggesting that *EIF4EBP1* expression can stratify patients within the most aggressive NB subset. We additionally analyzed the prognostic value of *EIF4EBP1* expression in stage 4 NB patients. We found high *EIF4EBP1* expression to significantly predict decreased overall and event-free survival of stage 4 patients in two independent cohorts (SEQC and NRC cohorts) (p=3.2e-04 Fig. 2g; p=3.8e-03, Fig. 2h and supplementary Fig. 1g&h). This highlights that *EIF4EBP1* expression robustly stratifies patients within the advanced NB subgroups. Altogether, our analyses support that *EIF4EBP1* expression is a factor of poor prognosis in all NB, as well as in high-risk and stage 4 NB.

**Figure 2:**
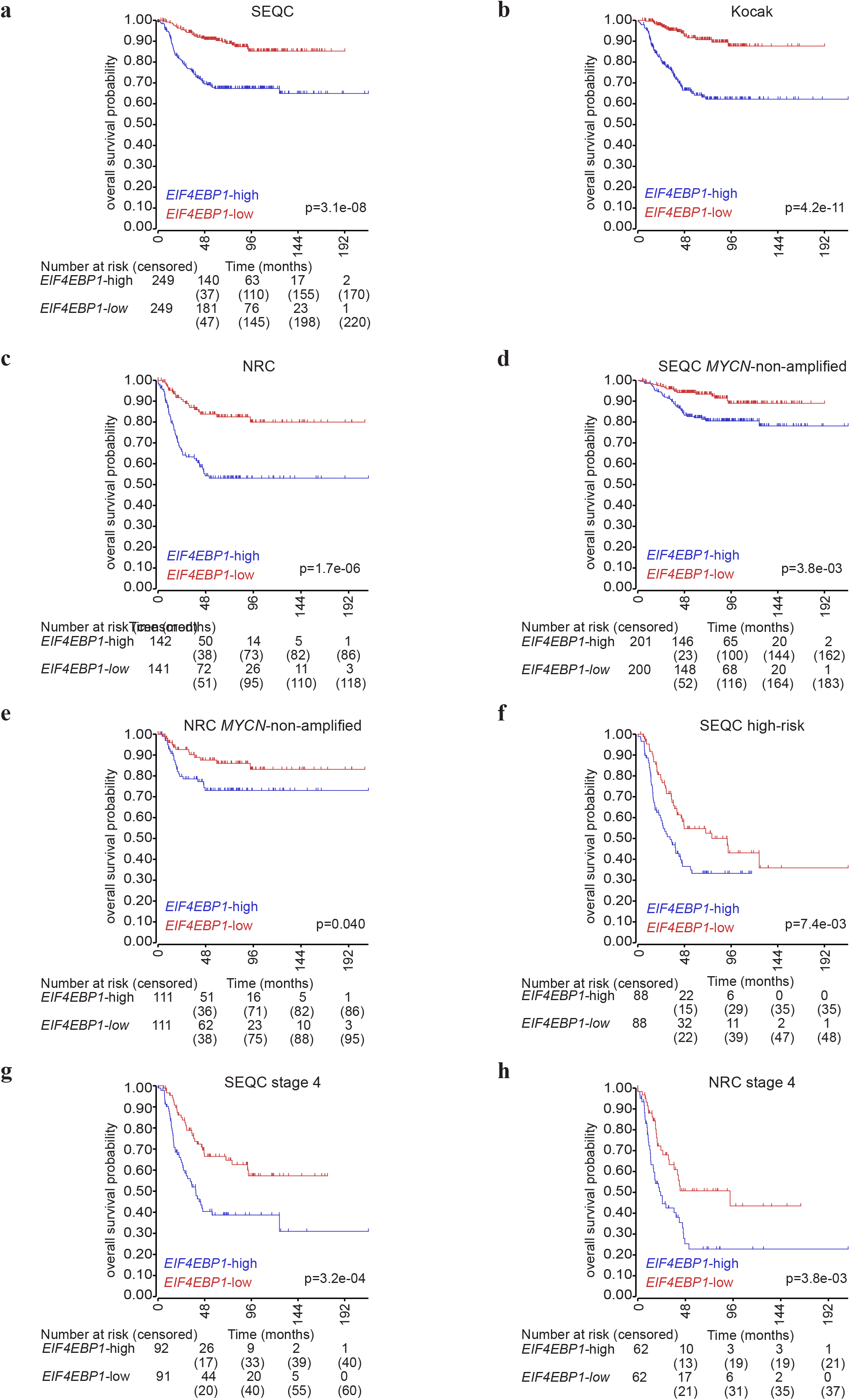
*EIF4EBP1* mRNA expression correlates with overall survival in NB patients. (a-c) Kaplan-Meier survival estimates of overall survival of NB patients stratified by their *EIF4EBP1* mRNA expression levels (median cut off) in the SEQC (a), Kocak (b) and NRC (c) cohorts. (d-h) Kaplan-Meier survival estimates of overall survival of patients with *MYCN*-non-amplified NB (d, e), high-risk NB (f) or stage 4 NB (g, h) stratified by their *EIF4EBP1* mRNA expression levels in the indicated NB cohorts. The number of patients at risk (or censored) are reported under the Kaplan-Meier plot by four-year intervals; such information were not accessible for the Kocak cohort. Significance was determined by log rank test. Data were obtained from the R2: Genomics Analysis and Visualization Platform.

**Table 1.** Multivariate analysis for overall survival of NB patients in the SEQC cohort.

| **Variables** | **HR** | **95.0% CI** | **p value** |
| --- | --- | --- | --- |
| *MYCN* amplification | 22.373 | 8.89-56.306 | 0 |
| High *EIF4EBP1* mRNA expression | 2.16 | 1.255-3.717 | 0.005 |
| *MYCN* amplification*high *EIF4EBP1* mRNA expression | 0.222 | 0.08-0.614 | 0.004 |
| **Variables** | **HR** | **95.0% CI** | **p value** |
| Stage 4 | 17.618 | 6.694-46.366 | 0 |
| High *EIF4EBP1* mRNA expression | 5.457 | 2.026-14.697 | 0.001 |
| Stage 4*high *EIF4EBP1* mRNA expression | 0.292 | 0.097-0.879 | 0.029 |
| **Variables** | **HR** | **95.0% CI** | **p value** |
| Age at diagnosis | 33.018 | 7.835-139.139 | 0 |
| High *EIF4EBP1* mRNA expression | 12.204 | 2.832-52.598 | 0.001 |
| Age at diagnosis*high *EIF4EBP1* mRNA expression | 0.16 | 0.035-0.74 | 0.019 |

**Table 2.** Multivariate analysis for overall survival of NB patients in the NRC cohort.

| Variables | HR | 95.0% CI | p value |
| --- | --- | --- | --- |
| *MYCN* amplification | 4.967 | 1.118-22.066 | 0.035 |
| High *EIF4EBP1* mRNA expression | 3.031 | 1.543-5.954 | 0.001 |
| *MYCN* amplification*high *EIF4EBP1* mRNA expression | 0.656 | 0.135-3.181 | 0.601 |
| Variables | HR | 95.0% CI | p value |
| Stage 4 | 15.050 | 4.239-53.432 | 0.018 |
| High *EIF4EBP1* mRNA expression | 5.144 | 1.330-19.895 | 0 |
| Stage 4*high *EIF4EBP1* mRNA expression | 0.36 | 0.081-1.598 | 0.179 |
| Variables | HR | 95.0% CI | p value |
| Age at diagnosis | 0.27 | 0.036-2.056 | 0 |
| High *EIF4EBP1* mRNA expression | 0.364 | 0.048-2.772 | 0.002 |
| Age at diagnosis*high *EIF4EBP1* mRNA expression | 55.427 | 3.258-942.88 | 0.005 |

### High 4EBP1 protein expression is associated with prognostically unfavorable histology of NB

To independently confirm the prognostic value of *EIF4EBP1*/4EBP1 in NB and to determine the biomarker potential of 4EBP1 protein expression in NB, we immunohistochemically analyzed NB TMAs consisting of 69 patient samples. Staining of the TMAs with a 4EBP1-specific antibody revealed a cytoplasmic staining (Fig. 3a), consistent with the expected cellular localization of 4EBP1 [43]. We semi-quantitatively evaluated 4EBP1 staining intensity and correlated 4EBP1 immunoreactivity with the NB histological subtypes according to the International Neuroblastoma Pathology Classification (INPC), which distinguishes patients with favorable or unfavorable histology based on grade of neuroblastic differentiation and mitosis-karyorrhexis index. We found that tumors with unfavorable histology more frequently exhibited a high 4EBP1 staining score (IRS 7-12) as compared to tumors with favorable histology (Fig. 3b), indicating that high 4EBP1 protein expression is associated with more aggressive NB subsets.

**Figure 3:**
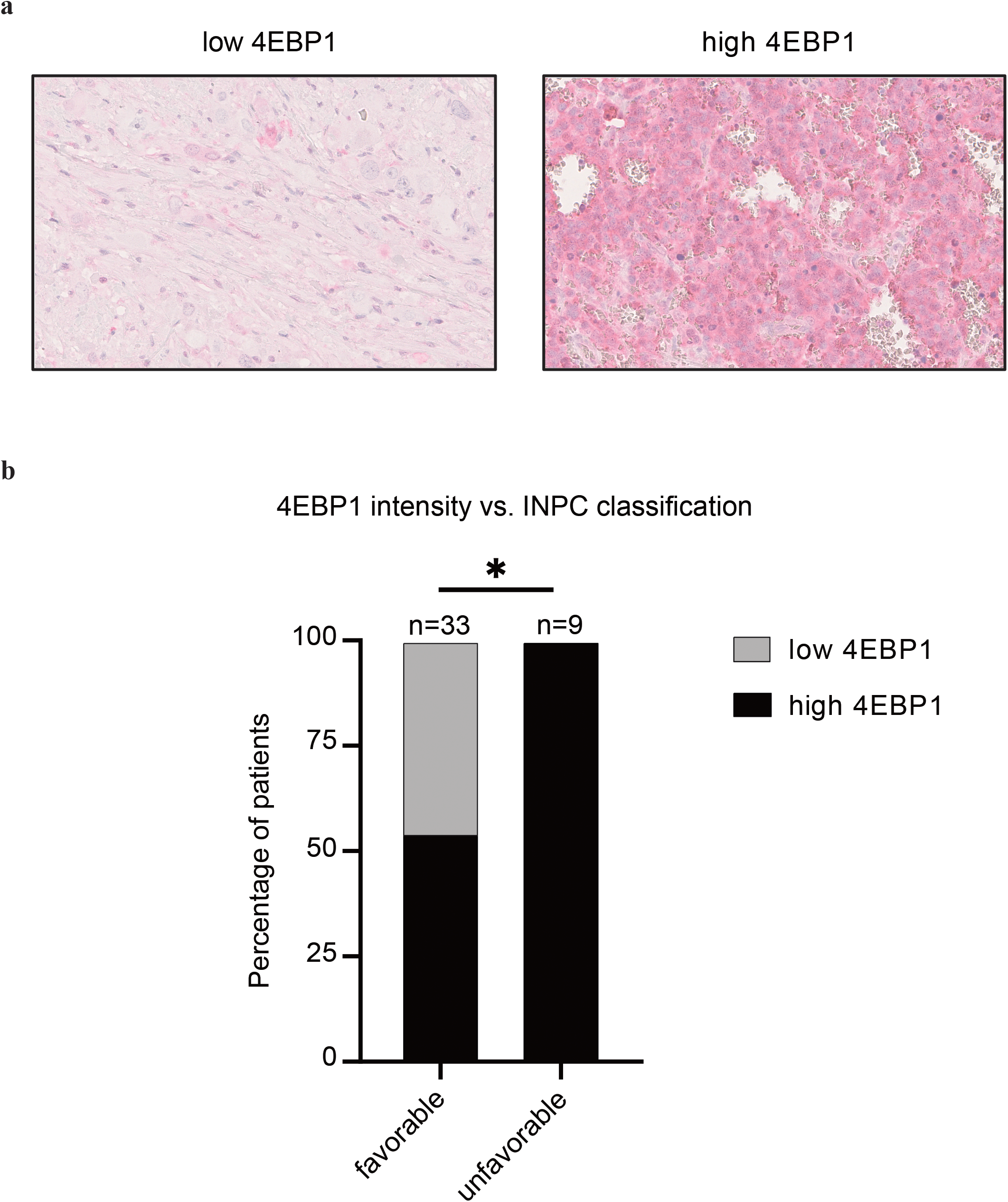
4EBP1 protein expression is associated with histological subtype of NB. (a) Representative images of low (left panel) and high (right panel) 4EBP1 immunohistochemical staining levels of selected NB samples represented on the NB TMAs. (b) Distribution of NB cases showing low (IRS 0-6) versus high (IRS 7-12) 4EBP1 protein expression in prognostically favorable versus unfavorable histological subtypes according to International Neuroblastoma Pathology Classification (INPC). Fisher’s exact test was used to calculate significance. \**P*<0.05.

### *EIF4EBP1* promoter activity and transcription is controlled by MYCN

To delineate how elevated *EIF4EBP1* expression is mechanistically connected to *MYCN* amplification and overexpression in NB, we investigated the transcriptional regulation of *EIF4EBP1* by MYCN. A previous report detected the presence of MYCN on *EIF4EBP1* promoter by ChIP in BE(2)-C, a *MYCN*-amplified NB cell line [38,39]. We validated and further extended this finding by analyzing ChIP-seq data available from an additional *MYCN*-amplified NB cell line, Kelly. This revealed that MYCN binds the endogenous *EIF4EBP1* promoter region (which encompasses exon 1 and a part of intron 1) at three distinct positions, indicating three potential MYCN binding sites (Fig. 4a). *In silico* analysis of the promoter region sequence confirmed the presence of structural E-boxes at the three occupied locations (Fig. 4b). To evaluate the impact of MYCN on the regulation of *EIF4EBP1* promoter activity, we designed a luciferase-based gene reporter assay by cloning the *EIF4EBP1* promoter region (−192 to +1372) in front of a Firefly Luciferase gene (Fig. 4b). The activity of the wildtype *EIF4EBP1* promoter was dose-dependently increased upon forced expression of MYCN in HEK-293-T cells (supplementary Fig. 2a), which was accompanied by an upregulation of endogenous 4EBP1 protein level (supplementary Fig. 2b). To investigate which E-boxes are necessary for the transcriptional activation of the *EIF4EBP1* promoter by MYCN, either a single or a combination of two of the three potential binding sites were mutated. Mutation of either of the three binding sites alone was sufficient to reduce MYCN-induced promoter activity by at least 50% (Fig. 4c). Any combinations of two mutated binding sites virtually abolished promoter activity driven by MYCN overexpression (Fig. 4c), suggesting that two binding sites, without a specific preference of one over another, are needed for full induction of *EIF4EBP1* promoter activity by MYCN. We next intended to confirm whether MYCN directly regulates *EIF4EBP1* transcription in NB cell lines. To do so, we chose two *MYCN*-amplified NB cell lines, IMR-32 and Kelly, in which we knocked down MYCN expression by siRNA and examined the impact on *EIF4EBP1* mRNA levels by qPCR. The depletion of *MYCN* caused a significant reduction of *EIF4EBP1* transcript levels in both cell lines (Fig. 4d&e). To further support these observations, we assessed the impact of forced MYCN expression on *EIF4EBP1* transcript and protein levels by using SHEP-TR-MYCN cells, which are *MYCN*-non-amplified NB cells engineered to express exogenous MYCN with a tetracycline inducible system [17]. Doxycycline treatment markedly increased *EIF4EBP1* mRNA level over time (Fig. 4f), in parallel with progressive upregulation of MYCN expression (Fig. 4g). This was accompanied by a net increase in the 4EBP1 protein level (Fig. 4g), supporting that MYCN positively controls *EIF4EBP1* mRNA and protein expression in NB cells. In accordance with these findings, analyses of expression data from a transgenic mouse model of MYCN-driven NB (TH-MYCN; [44]) revealed that *EIF4EBP1* expression is upregulated in NB tumors as compared to the corresponding normal tissue, i.e. the ganglia (Fig. 4h). Taken together, our data provide further evidence that *EIF4EBP1* is a transcriptional target of MYCN, potentially providing a mechanistic basis for the observed overexpression of *EIF4EBP1* in *MYCN*-amplified NB patients.

**Figure 4:**
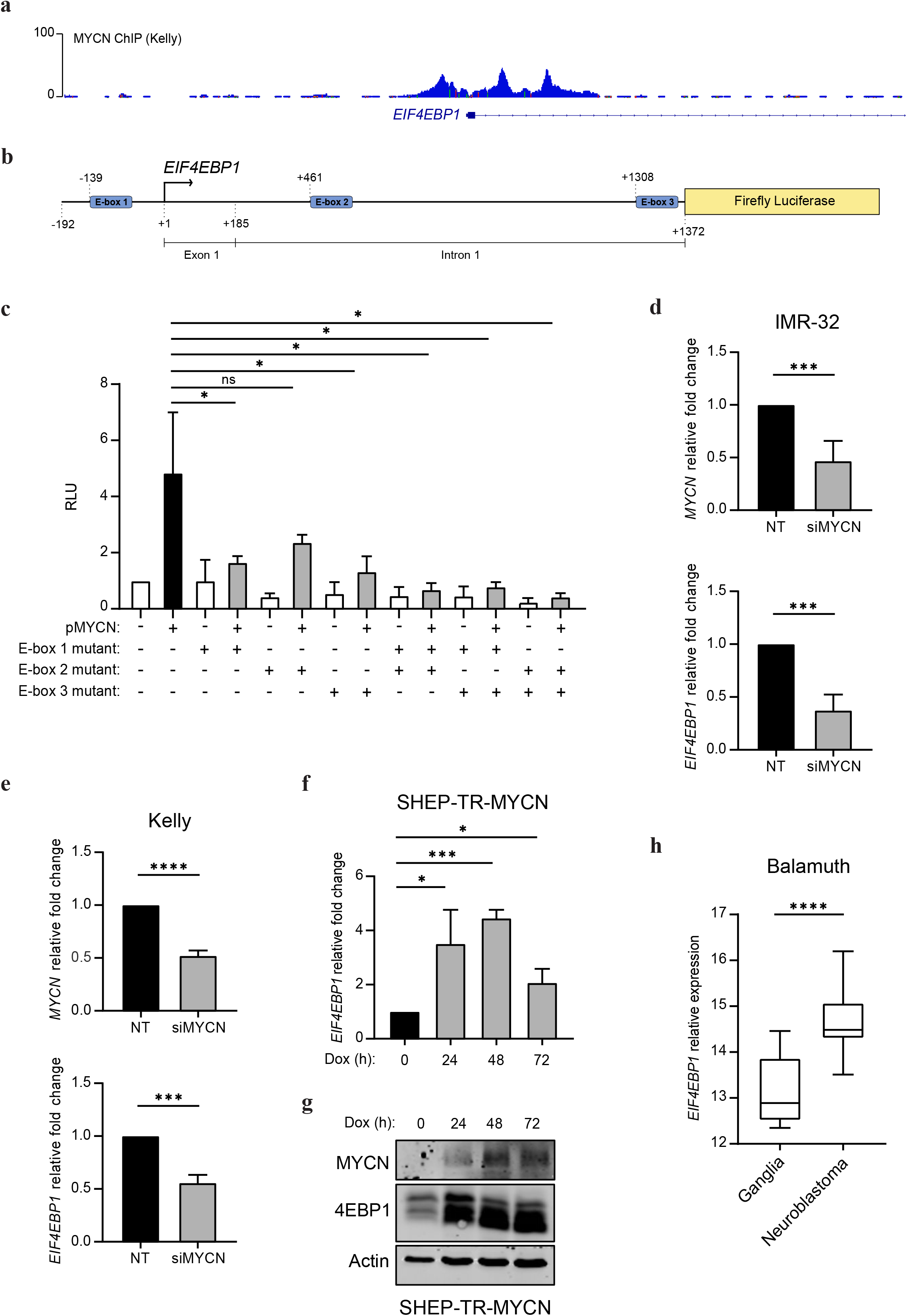
*EIF4EBP1* promoter activity and expression is regulated by MYCN. (a) ChIP peaks of MYCN in the *EIF4EBP1* promoter region in Kelly NB cell line. (b) Scheme of the *EIF4EBP1* promoter reporter highlighting the three E-boxes corresponding to MYCN binding sites. (c) HEK-293-T were transfected with wildtype or different E-box mutants *EIF4EBP1* promoter Firefly Luciferase constructs with or without a MYCN expressing plasmid (pMYCN). A *Renilla* Luciferase vector was used as an internal control. (d, e) Relative *MYCN* and *EIF4EBP1* mRNA levels upon siRNA-mediated knockdown of *MYCN* in the *MYCN*-amplified IMR-32 (d) and Kelly (e) cell lines, as measured by qRT-PCR. (f, g) SHEP-TR-MYCN cells were treated with doxycycline (1 μg/ml) for the indicated times; *EIF4EBP1* mRNA levels were determined by qRT-PCR (f) and levels of MYCN and 4EBP1 proteins were monitored by immunoblot using the indicated antibodies (g). (h) Relative *EIF4EBP1* mRNA expression in healthy control tissues (ganglia, n=9) and NB tumors (n=26) of a TH-MYCN transgenic mouse model of NB (Balamuth’s dataset; [44]). Data were retrieved from the R2: Genomics Analysis and Visualization Platform. Statistics were determined using Student’s t-test or Mann-Whitney U-test. Exact p-values are presented. \**P*<0.05, \*\**P*<0.01, \*\*\**P*<0.001, \*\*\*\**P*<0.0001.

## DISCUSSION

*MYCN*-amplification is accountable for aggressive NB subsets as it has been associated with increased risk of relapse and reduced overall survival of patients [6]. Since MYCN is considered “undruggable”, there is a demand for identifying targetable downstream effectors of MYCN [18,19]. In addition, since NB is a clinically heterogenous disease, ranging from spontaneous regression to progression despite aggressive therapies, novel markers that improve patient risk stratification and hence allow for optimal treatment allocation are warranted [4]. Here, we report that *EIF4EBP1* expression levels are significantly elevated in NB compared to corresponding non-tumor tissues and positively correlate with both *MYCN* expression and *MYCN* amplification status in at least two independent NB patient cohorts. Furthermore, using three independent NB cohorts, we report that high *EIF4EBP1* expression is a strong predictor of poor overall and event-free survival across all NB patients. This was not independent of *MYCN* amplification status, tumor stage or age at diagnosis, which can be explained in part by the regulation of *EIF4EBP1* promoter by MYCN which we characterized. However, *EIF4EBP1* expression can predict prognosis within distinct patient groups like the *MYCN*-non-amplified patients subset, for which little biomarkers have been identified. Moreover, we observed that high *EIF4EBP1* expression was associated with poor prognosis in the group of patients with aggressive stage 4 NB. Of note, less than a third of stage 4 patients carry a *MYCN* amplification. Thus, it may be worth considering that, in addition to *MYCN* amplification status, levels of *EIF4EBP1* expression could help identifying patients carrying clinically more aggressive tumors within the stage 4 NB patients group. *EIF4EBP1* expression was also linked to worse outcome among high-risk NB patients. Given that *MYCN* amplification is not able of predicting outcome within high-risk NB patients [45], it appears that *EIF4EBP1* expression has a prognostic power beyond *MYCN* amplification in this patient subset. Thus, *EIF4EBP1* expression may represent a promising biomarker for prognostic stratification of high-risk NB patients, in addition to the recently reported genetic alterations in the RAS and p53 pathways [11]. This is further supported by the association we observed between high 4EBP1 protein expression and unfavorable NB histological subtype. Together, our findings highlight a previously underappreciated prognostic factor, i.e., *EIF4EBP1*/4EBP1, which may help refining risk stratification of NB patients, including *MYCN*-non-amplified, stage 4 and high-risk patients, and could potentially assist in tailoring more personalized treatment options. Beyond NB, *EIF4EBP1* expression was reported to be a factor of poor prognosis in breast and liver cancers [28,31], as well as in all TCGA tumor types combined [27]. While our data indicate that *EIF4EBP1* expression has prognostic power in pediatric cancer, together this supports that *EIF4EBP1* expression represents a factor of poor prognosis in a large number of different tumor types.

Our study also extends previous knowledge by providing further experimental evidence to explain the association between *EIF4EBP1* and *MYCN* expression in NB and the overexpression of *EIF4EBP1* in *MYCN*-amplified NB. Our data revealed that MYCN induces transcription of *EIF4EBP1* by regulating its promoter through multiple binding sites, which was originally suggested by detection of MYCN binding to the *EIF4EBP1* promoter by ChIP analysis [38,39]. However, whether MYCN could transcriptionally regulate the *EIF4EBP1* promoter was still elusive. We demonstrate that MYCN activates the *EIF4EBP1* promoter through binding at three distinct E-boxes, which in turn leads to transcriptional increase of *EIF4EBP1*. Together with the previous ChIP analysis, this supports that *EIF4EBP1* is a direct target gene of MYCN in NB cells. These findings are in line with a previous study reporting that MYC controls *EIF4EBP1* by binding its endogenous promoter in colorectal cancer cells [46], highlighting a general regulation of *EIF4EBP1* by MYC family members in cancer cells. Expression levels of *EIF4EBP1* appear not only elevated in *MYCN*-amplified versus *MYCN*-non-amplified NB but are also upregulated in *MYCN*-non-amplified tumors relative to control tissue. It might be speculated that in *MYCN*-non-amplified NB, *EIF4EBP1* expression may be regulated by transcription factors other than MYCN. In particular, ATF4, which is critical for the metabolic response of NB cells to glutamine starvation [47,48], has been shown to control *EIF4EBP1* promoter and transcription in pancreatic beta cells [36]. This transcription factor is highly expressed in NB, and in particular in advanced stage 4 [48]. In addition, another transcription factor that is commonly overexpressed in NB is OCT4 [49]. Of note, this transcription factor has been identified by ChIP-seq to bind the promoter region of *EIF4EBP1* in human embryonic stem cells [50,51], thus OCT4 may also activate *EIF4EBP1* transcription in NB cells. Together, these data suggest potential mechanisms underlying the MYCN independent regulation of *EIF4EBP1* expression in *MYCN*-non-amplified NB patients. Given the prognostic significance of *EIF4EBP1*/4EBP1 in NB, it is possible that 4EBP1 confers advantages to NB tumor growth or tumor cell survival. As evidenced by the presence of necrotic areas flanked by HIF-1α positive staining [52], NB experience metabolic stress, corresponding to nutrient deprivation and hypoxia, as a consequence of abnormal and immature vascularization [53,54]. One important mechanism for cancer cells to adapt to metabolic stress is through reprogramming of mRNA translation [55]. As a major regulator of mRNA translation, 4EBP1 may aid NB cells to cope with hypoxia and nutrient deprivation. This is supported by the report that 4EBP1 promotes survival of breast tumors under hypoxia by stimulating the synthesis of pro-angiogenic factors, like HIF-1α and VEGF, to facilitate tumor angiogenesis *in vivo* [26]. In addition, the control of mRNA translation was shown to be critical to prevent the deleterious effects of MYCN and MYC overexpression, as we and others previously reported [46, 56]. In fact, 4EBP1, by reducing overall protein synthesis, was reported to prevent cell death induced upon MYC overexpression, likely by blunting accumulation of misfolded proteins and proteotoxic ER stress [46]. It is possible that in a similar manner 4EBP1 contributes to inhibit cell death induced by MYCN overexpression in *MYCN*-amplified NB.

In summary, the findings reported here indicate that *EIF4EBP1* is a direct target gene of MYCN in NB, explaining the observed high expression of *EIF4EBP1* in NB, and that *EIF4EBP1* mRNA and protein expression have prognostic values in NB patients, especially for stratifying high-risk NB patients.

## AUTHORS CONTRIBUTION

Conception and design: Kai Voeltzke and Gabriel Leprivier.

Provision of study material and patients: Irene Esposito and Thomas Kirchner.

Financial and administrative support: Guido Reifenberger.

Data analysis and interpretation: Kai Voeltzke, Thomas G. P. Grünewald, Alexander Schramm and Gabriel Leprivier.

Critical review and discussion: Barak Rotblat, Marc Remke, Alexander Schramm, Guido Reifenberger and Gabriel Leprivier.

Experimental support: Kai Voeltzke, Katerina Scharov, Cornelius Funk, Alisa Kahler, Daniel Picard, Laura Hauffe and Martin F. Orth.

Manuscript writing: Kai Voeltzke, Guido Reifenberger and Gabriel Leprivier. Final approval of the manuscript: All authors.

## ACKOWLEDGMENTS

We would like to thank Dr. Bastian Malzkorn (Institute of Neuropathology, Heinrich Heine University Düsseldorf) for helpful discussions.

## DATA SHARING STATEMENT

The data that support the findings of this study are available from the corresponding author upon reasonable request.

## SUPPLEMENTARY FIGURE LEGENDS

**Supplementary Figure 1:**
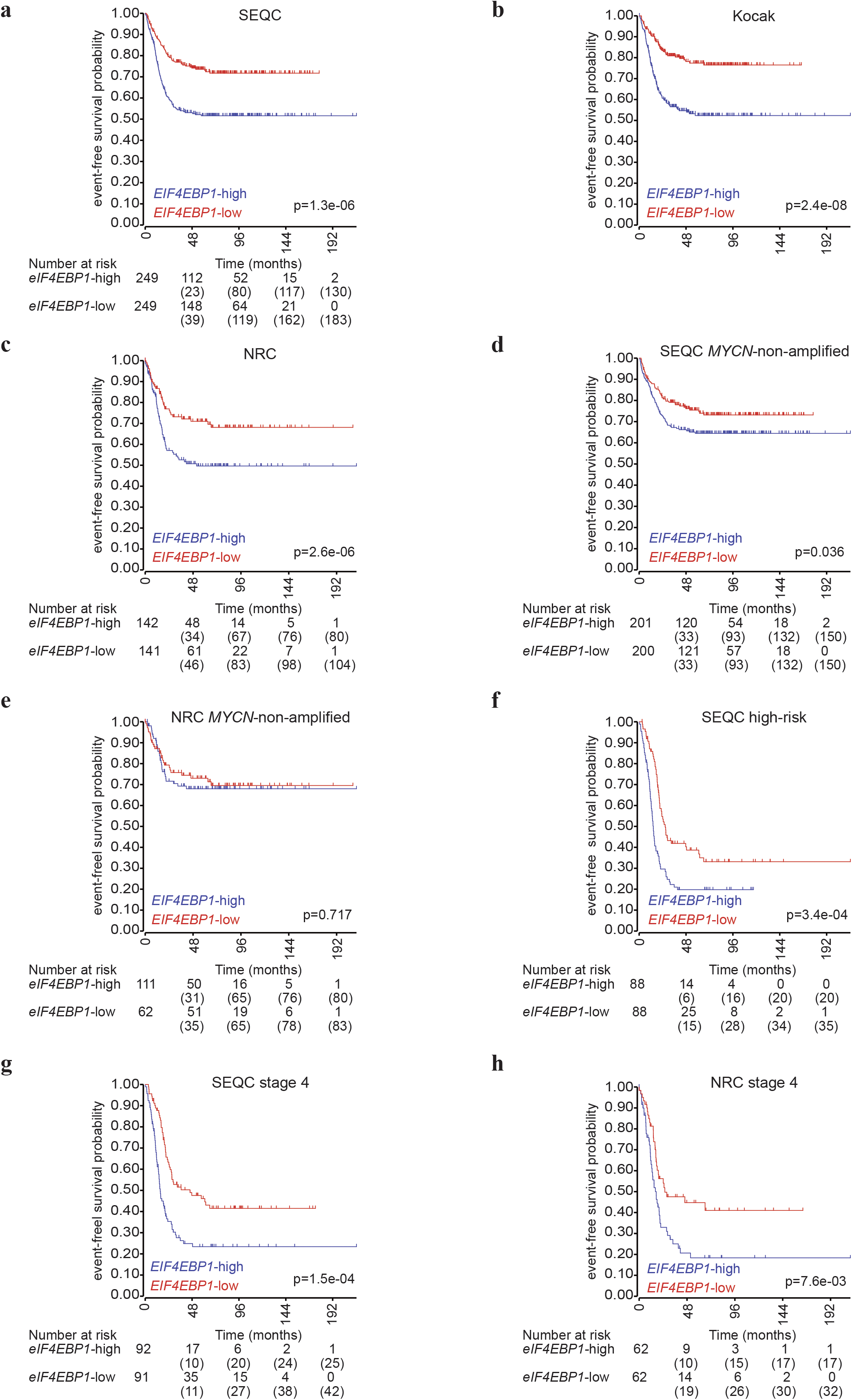
*EIF4EBP1* mRNA expression correlates with event-free survival in NB patients. (a-b) Kaplan-Meier survival estimates of event-free survival of NB patients stratified by their *EIF4EBP1* mRNA expression levels (median cut off) in the SEQC (a), Kocak (b) and NRC (c) cohorts. (d-h) Kaplan-Meier estimates of event-free survival of patients with *MYCN*-non-amplified NB (d, e), high-risk NB (f) or stage 4 NB (g, h) stratified by their *EIF4EBP1* mRNA expression levels in the indicated NB cohorts. The number of patients at risk (or censored) are reported under the Kaplan-Meier plot by four-year intervals; such information were not accessible for the Kocak cohort. Significance was determined by log rank test. Data were obtained from the R2: Genomics Analysis and Visualization Platform.

**Supplementary Figure 2:**
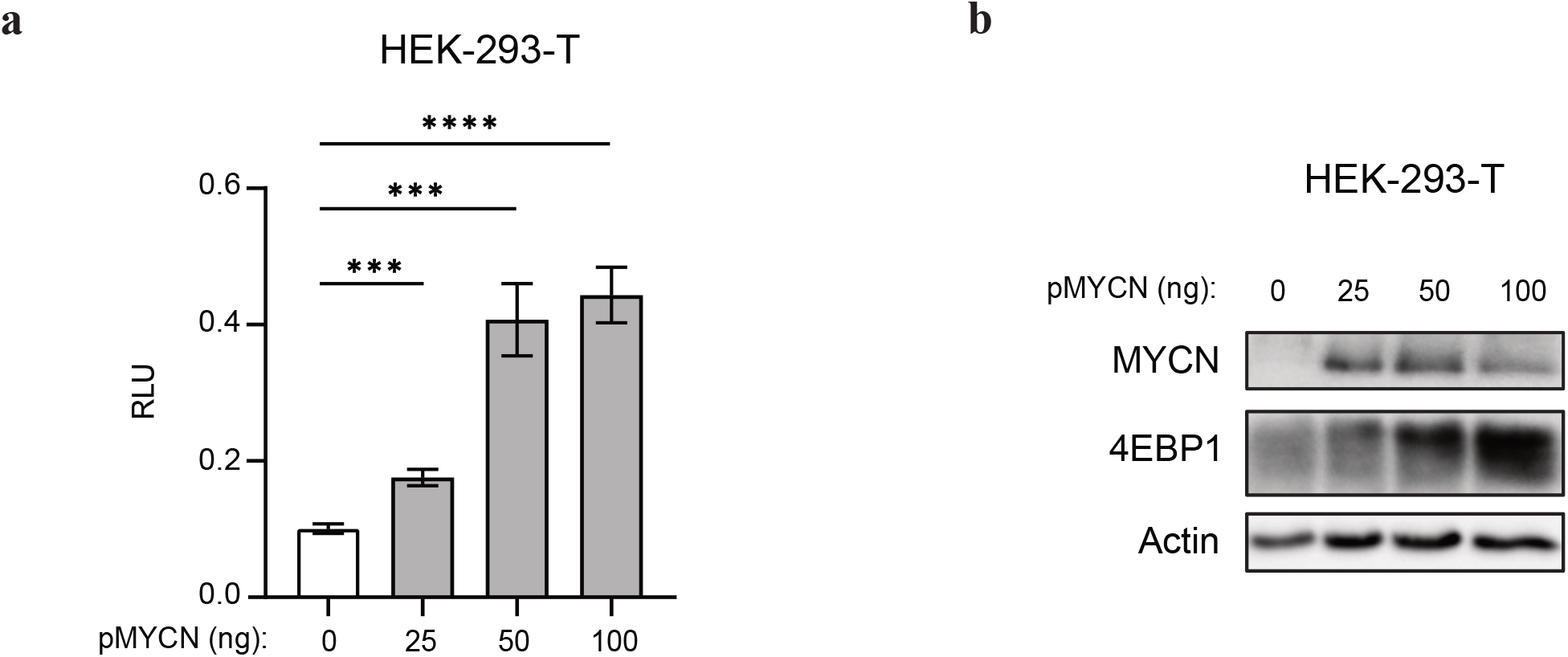
MYCN stimulates *EIF4EBP1* promoter activity and 4EBP1 protein expression in a dose-dependent manner. (a) HEK-293-T cells were transfected with the wildtype *EIF4EBP1* promoter Firefly Luciferase construct and with the indicated amounts of MYCN expressing plasmid (pMYCN). A *Renilla* Luciferase vector was used as an internal control. (b) MYCN and 4EBP1 protein expression was monitored in cell lysates from (a) by immunoblot analyses using the indicated antibodies.

**Supplementary table 1:**
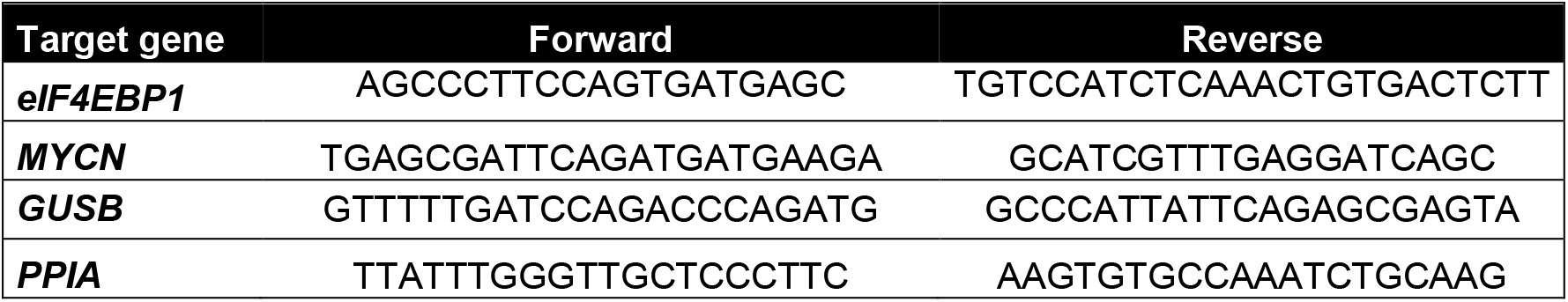
Primer list.

